# SINE Retrotransposons Import Polyadenylation Signals to 3’UTRs in Dog (*Canis familiaris*)

**DOI:** 10.1101/2020.11.30.405357

**Authors:** Jessica D. Choi, Lelani A. Del Pinto, Nathan B. Sutter

## Abstract

**Background:** Messenger RNA 3’ untranslated regions (3’UTRs) control many aspects of gene expression and determine where the transcript will terminate. The polyadenylation signal (PAS) AAUAAA is a key regulator of transcript termination and this hexamer, or a similar sequence, is very frequently found within 30 bp of 3’UTR ends. Short interspersed element (SINE) retrotransposons are found throughout genomes in high copy number. When inserted into genes they can disrupt expression, alter splicing, or cause nuclear retention of mRNAs. The genomes of the domestic dog and other carnivores carry hundreds of thousands Can-SINEs, a tRNA-related SINE with transcription termination potential. Because of this we asked whether Can-SINEs may help terminate transcript in some dog genes.

**Results:** Dog 3’UTRs have several peaks of AATAAA PAS frequency within 40 bp of the 3’UTR end, including four bp-interval peaks at 28, 32, and 36 bp from the end. The periodicity is partly explained by TAAA(n) repeats within Can-SINE AT-rich tails. While density of antisense-oriented Can-SINEs in 3’UTRs is fairly constant with distances from 3’end, sense-oriented Can-SINEs are common at the 3’end but nearly absent farther upstream. There are nine Can-SINE sub-types in the dog genome and the consensus sequence sense strands (head to tail) all carry at least three PASs while antisense strands usually have none. We annotated all repeat-masked Can-SINE copies in the Boxer reference genome and found that the young SINEC_Cf type has a mode of 15 bp for target site duplications (TSDs). We find that all Can-SINE types favor integration at TSDs beginning with A(4). The count of AATAAA PASs differs significantly between sense and antisense-oriented retrotransposons in transcripts. Can-SINEs near 3’UTR ends are very likely to carry AATAAA on the mRNA sense strand while those farther upstream are not. We also identified loci where Can-SINE insertion has truncated or altered a dog 3’UTR compared to the human ortholog.

**Conclusion:** Dog Can-SINE activity has imported AATAAA PASs into gene transcripts and led to alteration of 3’UTRs. AATAAA sequences are selectively removed from Can-SINEs in introns and upstream 3’UTR regions but are retained at the far downstream end of 3’UTRs, which we infer reflects their role as termination sequences for these transcripts.

## Background

Within genes the 3’ untranslated region (3’UTR) within the mRNA has many roles in regulating gene expression and post-translational changes to a protein [1]. Mutations in 3’UTRs can affect gene expression in part by creating multiple isoforms of the gene, which is common in vertebrates [2]. In humans, 51-79% of genes express alternative 3’UTRs [2]. In pigs, a SNP in the 3’UTR of *CDC42* affects the mRNA expression in placentas by altering their binding affinity [3]. In another case, a SNP in the 3’UTR of *FNDC5* influences the mRNA stability in humans [4]. A pair of inverted *Alu* insertions in a reporter gene’s 3’UTR causes repression [5].

The median 3’UTR length in humans is approximately 1200 bp, while the median length in worms is only ~140 bp [6]. In vertebrates the longer 3’UTRs contain many cis-regulatory sequences. For example, genes with alternative 3’UTRs can be regulated by elements contained in the 3’UTR [2]. The length of 3’UTRs is also dependent on the tissue: genes with tissue-restricted expression tend to have just a single 3’UTR while genes transcribed in many tissues tend to have multiple 3’UTRs [7]. Despite the evolutionary conservation of gene sequences in the lineages of vertebrates there are large changes in 3’UTR complexity coincident with increasing organismal complexity of tissues and cellular differentiation in organisms [8]. The expansion of the 3’UTR sequence during evolution enables increasing complexity of proteins and their transcripts [8]. AU-rich elements found within 3’UTRs are important cis-regulators [9,10]. Depending on what effector proteins are recruited, the mRNA can either be stabilized or destabilized, and the 3’UTR can have different functional outcomes [8,9]. For example, tristetrapolin is involved in AU-rich element decay [11]. Other factors are used in order to regulate mRNA decay, such as Hsp27 and Hsp70 [12,13].

A gene’s 3’UTR is used to signal the end of transcription [8]. The sequence AAUAAA, or a closely related hexamer, is termed the polyadenylation signal (PAS) and is a key regulator of termination and polyadenylation [14]. AAUAAA is bound by a ternary complex of cleavage and polyadenylation specificity factor-160, WDR33, and CPSF-30 [15,16]. There are also other motifs that help provide the context for signaling the end of transcription. From the upstream end down they are: U-rich region (DNA: TGTA/TATA), the PAS, another U-rich region, a GTGT/GTCT, and a G-rich region [17].

Although AAUAAA (AATAAA in the DNA sense strand) is the most commonly used PAS, Beaudoing et al [18] identified 10 other variants, each just one mutation step away from AATAAA, that also have an increased frequency in the 3’ ends of 3’UTRs, albeit mostly with much smaller enrichment signals than AATAAA [18,19]. AATAAA and ATTAAA PASs are the most frequently used hexamers, seen in 58.2% and 14.9% of the 3’ fragments studied [18]. The commonest distance for both is −15 / −16 bp upstream of the 3’ end of the 3’UTR (measured as the distance from the nearest end of the PAS).

PASs closest to the 3’ end of the 3’UTR tend to be more conserved than those upstream in introns [20]. Older genes with a longer evolutionary history are more likely to have alternative polyadenylation sites [20,21]. In eight ENCODE human cell lines, 9.4% of termination signals occur within transposable elements (TEs) [22] and termination signals farther upstream are more often associated with transposable elements than are the canonical PASs [20]. Thus, transposable element insertions into gene transcripts can provide alternate polyadenylation signals. Human PASs not conserved in the mouse ortholog are associated with TEs; ~94% of human sites that are TE-associated are not conserved in the mouse [20]. TE insertion can alter transcript termination.

Gene expression in general is profoundly impacted by retrotransposons. Retrotransposon insertions are found throughout the genomes of vertebrates in the form of short interspersed elements (SINEs) and long interspersed elements (LINEs) [23]. SINEs and LINEs use a copy-paste method for insertion using an RNA intermediate (Pol III for SINEs, Pol II for LINEs). They have contributed to the expansion of the genome and provide a “fossil” record of genome change useful for cladistics. Mammalian-wide interspersed repeats (MIRs) are found in all mammals and are considered a record of a major genetic event that occurred before the radiation of mammals [24]. LINEs are autonomous retrotransposons that, like SINEs, show a strong orientation bias within transcripts and are present at high densities in introns [25,26].

Retrotransposons affect gene expression through several distinct mechanisms. They can become an exon (“exonization”), alter mRNA splicing, or cause mRNA to be retained in the nucleus or destroyed when SINE or LINE pairs are close and inverted [27,28]. Inverted SINE repeats occur at low frequency in transcripts and intergenic sequences [25,29]. Retrotransposons also regulate mRNAs through the piRNA pathway’s repression of gene expression in 3’UTRs [30]. Primate *Alu* and rodent B2 SINEs were recently reported to have ribozyme activity in the context of epigenetic suppression of gene expression [31]. In humans, *Alu* elements within 3’UTRs can also cause A-to-I editing that impacts gene expression [32].

SINEs have arisen many times in evolution and each type generally has an identifiable homology to one of its host genome’s own RNA genes, sometimes 5S rRNA or 7SL but most often a tRNA [33]. The tRNA-like head region of many SINEs contains internal RNA pol III promoter elements, the A and B boxes [34]. SINE RNA transcripts utilize “partner” LINE-encoded proteins to enable retrotransposition into a new locus [35]. This insertion process creates a direct repeat of sequence, called the target site duplication (TSD), that flanks the SINE sequence and provides a useful means of identifying the insertion event [36].

Carnivora-specific SINEs (Can-SINEs), first identified in the dog genome [37], are tRNA-derived SINEs [38,39] present in hundreds of thousands of copies in carnivore genomes but absent outside the order [36,40]. Like other tRNA-derived SINEs, they contain A and B box internal promoter sites for RNA pol III transcription. Like other T+ category SINEs they contain sequence motifs with potential for signaling transcription termination [41] such as AATAAA, and the TCT_3-6_ region. This last motif is an RNA pol III transcription terminator sequence, and is also found, for example, in mouse B2 SINEs near the 3’ end [42]. Despite being RNA pol III transcribed, some SINE transcripts are polyadenylated and these have a longer half-life [43]. It is unknown whether Can-SINEs are polyadenylated but young insertions often have long homopolymer runs of A(n). Can-SINEs have been annotated in the genomes of both canoidea (“dog like”) and feliform carnivores [44] and in every genome have distinct abundance and age profiles as measured by sequence divergence from a consensus [39,45,46].

Dogs have nine sub-types of Can-SINEs. SINEC_Cf is the youngest and its insertions often have less than 5% sequence divergence from the consensus. This type is apparently still active today as many insertions are so recent they are polymorphic [25,36,47]. These polymorphic SINEC_Cf insertions have played an important role in trait diversification within dogs; they are either associated with or are the causal component of numerous dog traits/disorders: narcolepsy (insertion intronic near *HCTR2* exon), merle coat pattern (intron-exon boundary in *SILV/PMEL*), black and tan coat (inverted SINEC_Cf repeat in an *ASIP* intron), retinal degeneration (within coding exon in *STK38L*), centronuclear myopathy (within coding exon in *PTPLA*), and retinal atrophy (near exon in *FAM161A*) [27,48-52].

Given the high copy number of SINE insertions in the dog genome, plus the potential for transcription termination signaling, we wondered if Can-SINEs import transcription termination signals used by dog genes. Like many other SINEs, dog Can-SINEs exhibit a strong orientation bias within transcripts, occurring at much higher frequency in antisense rather than sense orientation within introns [25]. Selection against termination signals present in sense oriented SINEs could be at least partly responsible. It has been known for over three decades that retrotransposons such as mouse B2 can provide polyadenylation signals to protein-coding genes [53]. In rabbits, CSINEs can provide functional polyadenylation signals in 3’UTRs [54]. Both B2 and CSINEs are T+ type SINEs like Can-SINEs. Retrotransposons inserted in 3’UTRs can also cause premature termination when they contain PASs. Insertion of a sense-oriented B2 SINE in the mouse *TNF* 3’UTR provides a PAS that shortens the 3’UTR by 650 nucleotides. This removes AU-rich elements further downstream in the 3’UTR, which in turn leads to overexpression [55]. In humans, *Alu*-PASs can produce a shortened transcript and some *Alu* elements also serve as microRNA targets [56,57]. When AATAAAs occur in *Alus*, they’re the result of single base mutations in the middle and ends of the *Alu* sequence. These *Alu*s used as major PASs are in sense orientation in 99% of cases [56,58].

We hypothesize that dog Can-SINEs have imported AATAAAs to genes that use them as PASs in 3’UTRs. Here we analyze the orientation bias of Can-SINEs in transcripts and the changing counts of AATAAAs within SINEs when the SINE is inserted within 3’UTRs vs. introns or intergenic sequences.

## Methods

### Obtaining Dog CanFam3.1 Reference Genome Datasets

We opted to use the CanFam3.1 reference genome of the dog although the CanFam4 reference became available during our work. We downloaded and used from the UCSC Genome Browser Table Browser the genome-wide Ensembl gene models (version 99), gaps, repeat track SINEs and LINEs, and chromosome start and end bp values. Gene models without an open reading frame (e.g. RNA genes) were excluded from the dataset because they are annotated as “UTR” and if left in would confuse signals relevant for 3’UTRs of protein-coding genes. From NCBI we downloaded the assembled chromosome fastas for the autosomes and chrX. These were used to programmatically slice out sub-sequences containing SINEs and their flanks as well as 3’UTRs.

### Defining Gene Features

With custom scripts we defined segments within genes such that 3’UTRs could be redundant but all bases within introns could not be part of any coding or untranslated exon segments. Intergenic sequence was defined as all sequences not present within gene model transcripts and not within the 500 bp upstream of transcription start sites.

### Analysis of Gene Features with Retrotransposons and Motif Finding

We wrote a Python script to count intersections of chromosome segment objects, find motifs such as AATAAA, and track strand orientations.

### Annotation of Can-SINEs

We downloaded the positions for all SINEC repeat-masked SINEs from the table browser of the UCSC genome browser reference for dog, CanFam3.1. A custom Python script retrieved the sequence plus 100 bp flanks for each SINE and attempted to identify perfectly matching target site duplications (TSDs) at a distance apart ranging from 1.25 to 0.75 times the length of the consensus for the given SINEC type. The longest perfect TSD found was kept and the minimum length of 6 bp was the filter for keeping a SINE copy in the analysis. The SINE was taken to be the sequence contained within the TSDs. A and B boxes were searched for within +/-10 bp of their position within the consensus for the SINEC type and up to 5 mismatches were permitted to still score a hit for box finding. The lowest mismatch box (or boxes) was kept. The longest A(n) string and longest string of A or T was identified. We also found the longest C or T string having at most 5 inclusions of non (C or T).

### Creation of sequence logos

We uploaded up to 10,000 TSD sequences for determining the sequence logos for Can-SINEs and up to 5000 A and 5000 B boxes sequences for their sequence logo determinations.

## Results

With an aim to better understand possible roles for Can-SINEs in dog 3’UTRs, we first confirmed the frequent presence of the AATAAA (AAUAAA in the RNA transcript) PAS near the 3’ end of 3’UTRs. We calculated the proportion of AATAAA and ATTAAA motifs found in sense orientation at each position for all dog 3’UTRs in the reference genome (Fig. 1). Our peak matches the −15 bp peak reported in Beaudoing et al [18] since we count distances from the 5’ end of each hexamer to the 3’UTR 3’ end, while they counted from the 3’ ends of hexamers. In addition to the overall peak with mode ~22-24 bp, AATAAA is found at high frequency at specific 4 bp intervals at 28, 32, and 36 bp from the 3’ end. The ATTAAA alternate PAS shows only a very small increase in frequency and the increases at ~22 bp in the other nine Beaudoing hexamers are not detectable in dogs (Table S1).

**Fig. 1.**
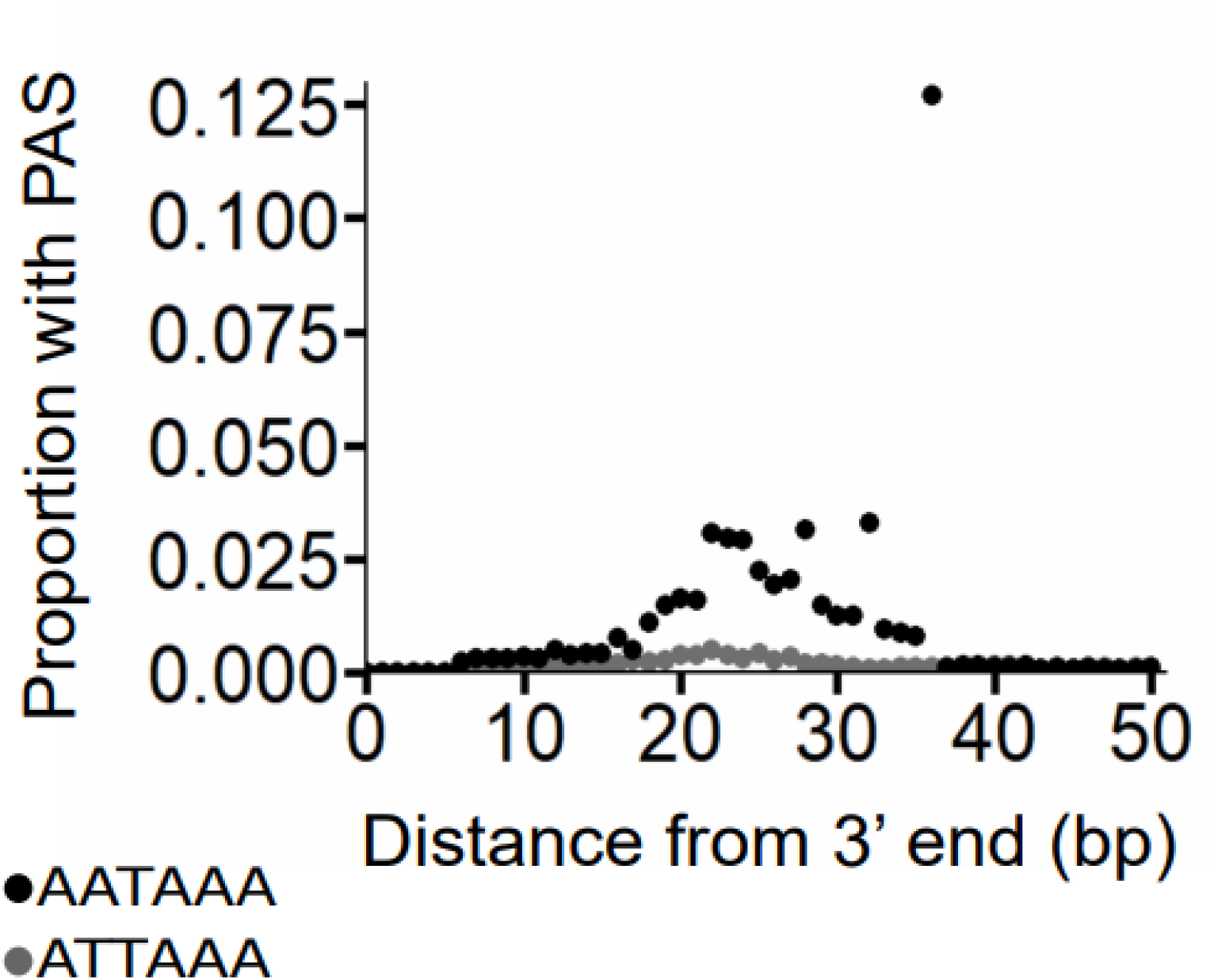
Dog 3’UTRs have multiple spikes in “AATAAA” PAS frequency. The distance to the 3’ end of the 3’UTR is calculated from the 5’-most base in each PAS hexamer.

SINE and LINE retrotransposons are inserted at high copy numbers throughout the dog genome, just as they are in the genomes of other species. They are present only at very low density in protein-coding exons and intronic sequences very near coding exons but in contrast are found at high density in introns, where they have an orientation bias in favor of antisense orientation [25,60]. We found that Can-SINEs and LINEs are also more prevalent in an antisense orientation in dog 3’UTRs providing the region analyzed is greater than 250 bp upstream of the 3’ end of the 3’UTR. LINEs have a strong antisense orientation bias in the region ~600-1000 bp upstream of 3’UTR end (Fig. 2b). In the last 250 bp of dog UTRs, in contrast, the LINE orientation bias disappears.

**Fig. 2.**
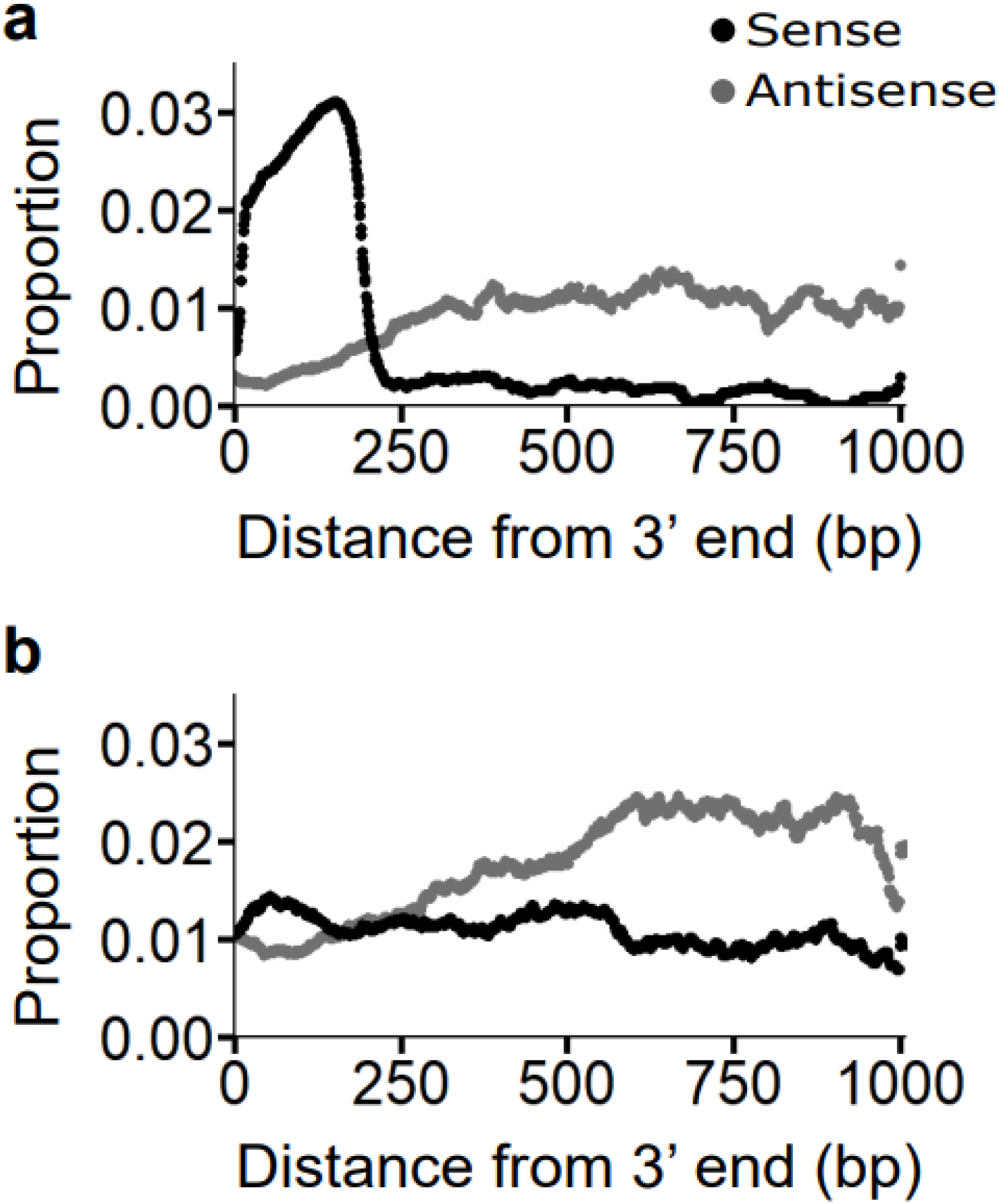
Density at the 3’ end of dog 3’UTRs for **a** Can-SINEs and **b** LINEs. At each base position, counting from the 3’ end, the proportion of all 3’UTRs having a base with that retrotransposon orientation is indicated.

We also assessed the density pattern for Can-SINEs by lumping together all the SINEC sub-types. This grouping includes older SINE types and a young type, SINEC_Cf, in which inserted elements have lower mean sequence divergence from the consensus. The Can-SINE density curve in 3’UTRs is X-shaped, with a strong reversal in orientation bias occurring at approximately 225-250 bp upstream of 3’UTR ends. While possibly coincidental, this is close to the length of a full-sized Can-SINE element. At the peak of the orientation bias, sense-oriented Can-SINEs are up to six times more frequent than antisense at the ends of dog 3’UTRs. Sense-oriented Can-SINEs are relatively frequently found at 3’ ends of 3’UTRs but are only rarely found farther than 250 bp upstream (Fig. 2a). We therefore hypothesized that Can-SINEs are active sequences in the context of 3’UTR ends.

In order to explore a relationship between SINEs and the PASs found at 3’UTR ends, we enumerated the number of PAS hexamers found in consensus sequences for Can-SINEs and MIRs in the dog genome (Fig. 3 and Table S2). When counting on the sense strand from SINE head to tail, all Can-SINE consensus sequences contain at least three polyadenylation signals clustered in their AT-rich tails. These tails typically contain some variant of AATA(n). SINEC_Cf, the youngest Can-SINE type, has four PASs within its tail. Dog Can-SINEs have a polypyrimidine repeat as well as the PAS motifs. The SINEC_Cf2, a2, b1, b2, and c2 consensus sequences also have a TCTTT motif for RNA pol III termination that directly follows the PAS. These sequences place Can-SINEs in the T+ category of SINEs alongside other mammal SINEs [38,39]. Notably, among the Can-SINE sub-types only SINEC_c1 and SINEC_c2 contain any antisense-oriented PASs in their consensus sequences and in both cases the hexamer, which is located in the head of the SINE at ~41 bp, is not an AATAAA. Thus, the sense vs. antisense strands of dog Can-SINEs may have starkly different potential to signal termination of RNA polymerase. MIR sub-types “MIR” and “MIRb” contain multiple PAS hexamer sequences in the consensus sense strand, however only one is AATAAA. MIR3 and MIRc do not contain PAS hexamers in the consensus sequences.

**Fig. 3.**
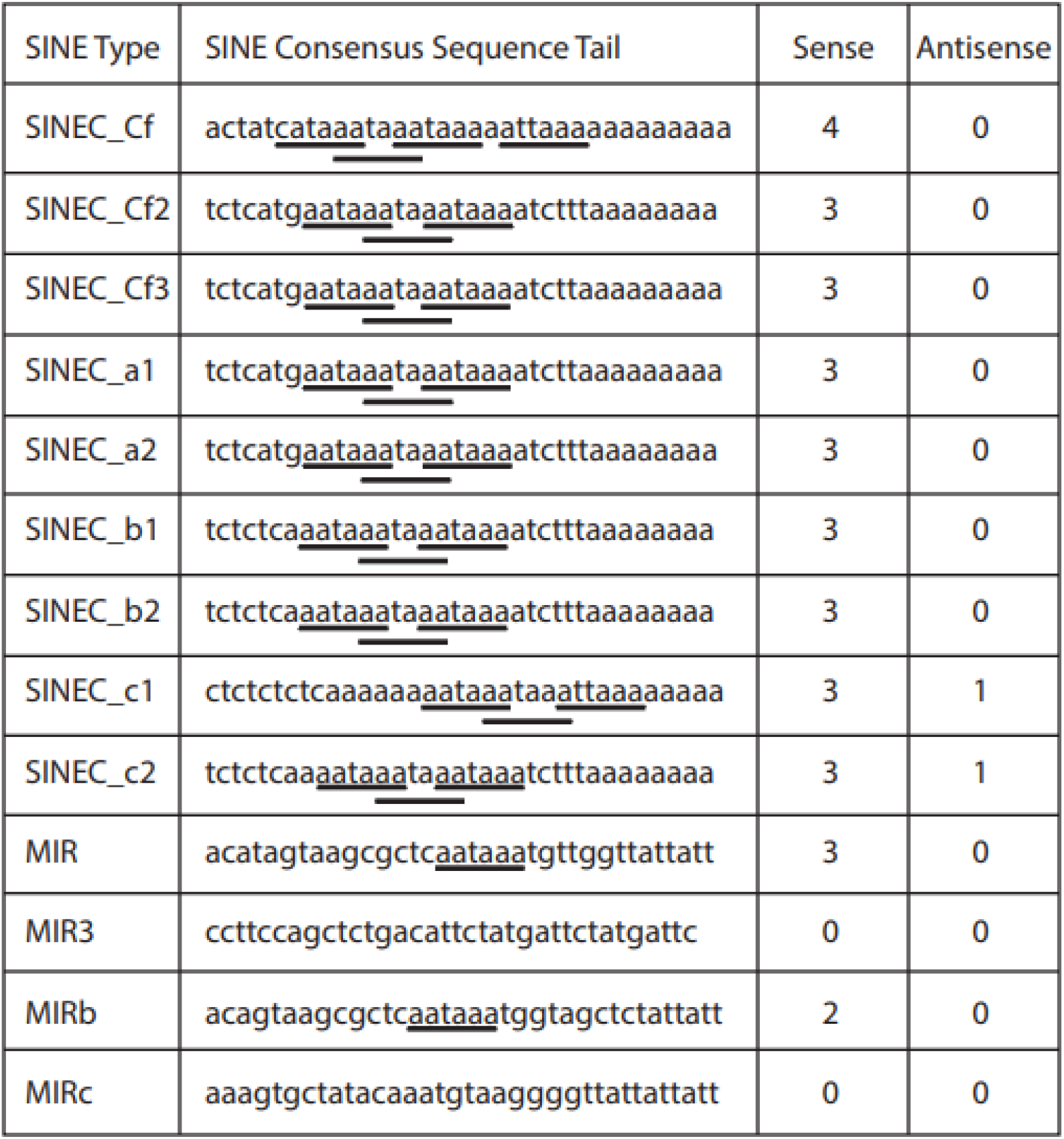
Can-SINE consensus sequences have multiple PASs in their sense (head to tail) strand. All 11 PAS sequences from Beaudoing et al, 2000, are included in the count. Counts cover the entire consensus sequence. Most Can-SINE PASs are “AATAAA” and are clustered in the A/T rich tail.

To better understand variation within the hundreds of thousands of individual SINE copies in the dog genome we collected all repeatMasked copies for the nine Can-SINE sub-types from the repeats track in the boxer reference genome CanFam3.1 (Fig. 4; Fig. S1). We analyzed each element’s sequence and flanks to identify Can-SINEs with perfect target site duplications. Of the 171,386 total SINEC_Cf copies in the reference, 124,901 (73%) have perfect TSDs between 10-20 bp (inclusive) in length. TSDs 15 bp long are the mode, with 26,055 SINEC_Cf copies (21%). This suggests that SINEC_Cf TSDs may be 15 bp long when they are made in the retrotransposition event that produces the new SINE copy.

**Fig. 4.**
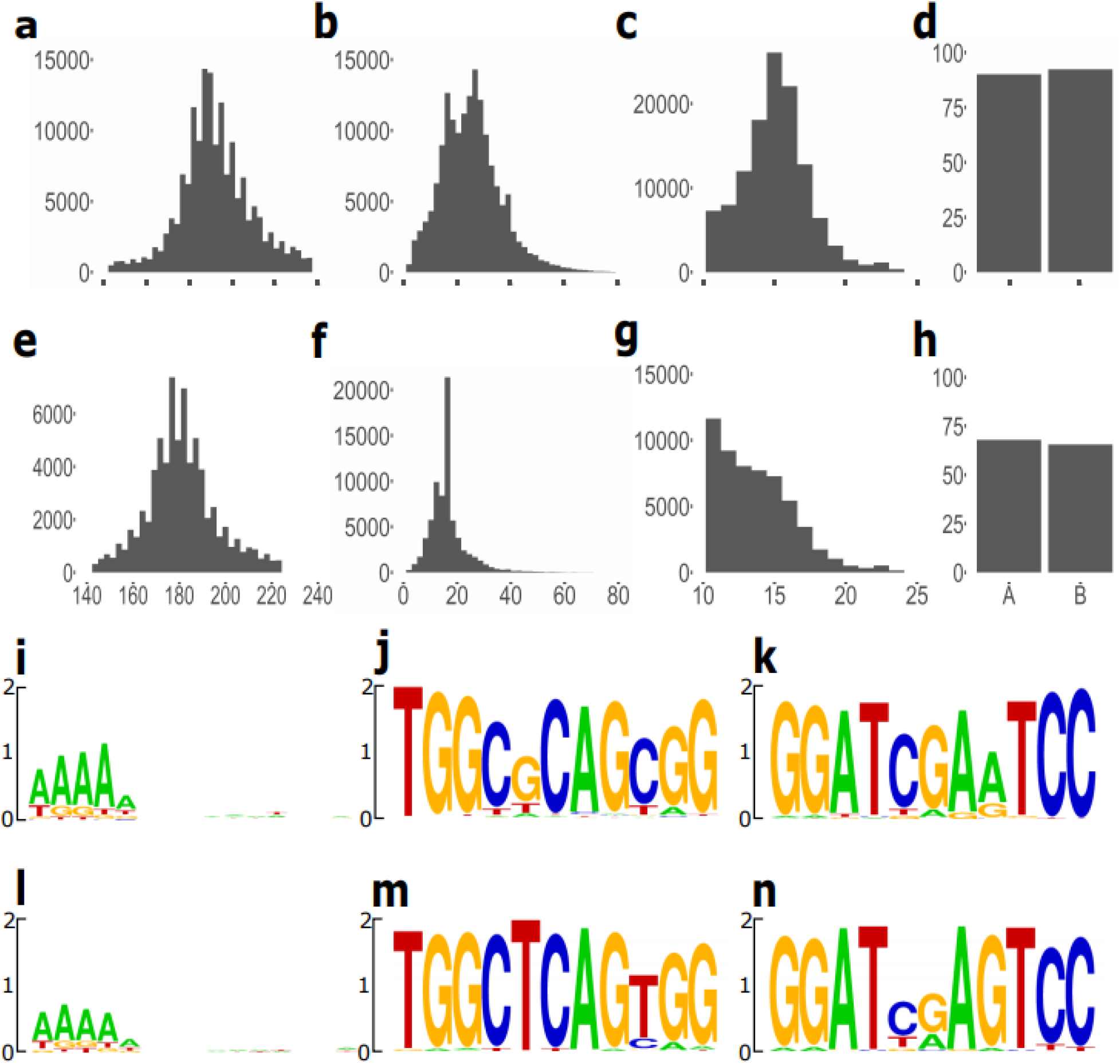
Annotation of Can-SINE sub-types SINEC_Cf (rows 1 and 3: **a-d, i-k**) and SINEC_a1 (rows 2 and 4: **e-h, l-n**). In panels **a-c** and **e-g** the y-axis is count of elements and the x-axis is bp. Length histograms of (**a, e**) the reference genome’s SINEs, (**b, f**) A/T-rich 3’ tails, (**c, g**) and target site duplications. (**d, h**) Barplots showing the percentage of SINE copies in which the A or B box internal promoter elements were identified. Compared to SINEC_a1 (row 2), SINEC_Cf (row 1) has a higher mean length, higher proportion of long perfect target site duplications, and a higher proportion of A and B boxes identified. (**i, l**) Sequence logos of target site duplications. Sequence logos of (**j, m**) A box and (**k, n**) B box.

In contrast to the relatively young SINEC_Cf type, other dog Can-SINEs are older. The elements in other dog Can-SINE types have higher mean divergence from consensus sequences and this can be readily seen, for example, in the TSD distribution for SINEC_a1 (Fig. 4g) where only a shoulder on the curve is evident at 15 bp. For most SINEC_a1 copies, one or both target sites have evidently mutated away from their original sequence. In fact, the differing ages of the Can-SINE types are apparent by comparing the TSD distributions in the relatively younger SINEC_Cf2 and SINEC_Cf3 (Fig. S1a, b), where perfect TSD peaks around 15 bp long are evident, vs. older types SINEC_c1 and SINEC_c2 that have very few 15 bp-long perfect TSDs (Fig. S1).

The SINEC_Cf copies with a perfect TSD 6 bp or greater (the “full set”) have a mean length inside the TSDs of 192 bp. This is the set used for Fig. 4a. SINEC_Cf’s with a perfect match 15 bp TSD have nearly identical mean length of 193 bp. The 15 bp TSD SINEC_Cfs have A and B boxes with Levenstein edit distances to consensus of 0.63 and 0.57, respectively, while in the full set these distances are greater, at 1.01 and 1.02 respectively. Plus, while 90% of SINE copies in the full set have a detectable A box and 92% have a detectable B box (Fig. 4d), these figures rise to 98% and 99%, respectively, for the 15 bp TSD set. Thus the 15-bp TSD set of SINEC_Cfs appears to skew slightly younger overall. Finally, the longest string of A or T (the tail) is an average of 25.1 bp long in the full set (Fig. 4b) but slightly longer, 27.2 bp, in the 15-bp TSD subset.

For all Can-SINEs in dog, the overriding sequence feature for the TSDs is an initial run of polyA (Fig. 4; Fig. S2). The A and B box sequences are very similar across the Can-SINE types, with, for example, just two positions varying in the A box between SINEC_Cf vs. SINEC_a1 (Fig. 4).

Given both the presence of PAS sequences in dog Can-SINE sense strands as well as the density patterns of Can-SINEs in 3’UTRs, we hypothesized that PASs in Can-SINEs are selected for or against depending on a particular Can-SINE’s orientation and position within gene transcripts. To assess this, we categorized Can-SINE insertions into four genomic locations (ignoring SINEs falling into other locations): intergenic, intronic (any position within an intron) or “near” or “far” from the 3’ end of a 3’UTR. To be categorized as “near” a Can-SINE must occur entirely within 225 bp of the 3’end of the 3’UTR. “Far” Can-SINEs are wholly contained within a 3’UTR and no closer to the 3’ end than 225 bp. Thus, some Can-SINEs spanning over 225 bp were excluded from this analysis. We categorized LINEs and MIRs in the same way and tracked retrotransposon orientation relative to the gene transcript. We then counted the number of AATAAA sequences on the head-to-tail strand of each retrotransposon and looked for differences in count distribution according to whether the retrotransposon was sense (RNA pol encounters the SINE head first) or antisense to the gene transcript. We find that the counts of AATAAAs are significantly different for some locations (Fig. 5). First, in intergenic sequences Can-SINEs, LINEs, and MIRs have the same mean AATAAA count in both orientations. In contrast, when inserted in introns, all three types of retrotransposons have lower mean AATAAA counts on their head-to-tail strands when the retrotransposon is inserted in sense orientation. Although the magnitude of the differences in mean are not great, there are >100,000 elements in each category and the differences in AATAAA count are significant.

**Fig. 5.**
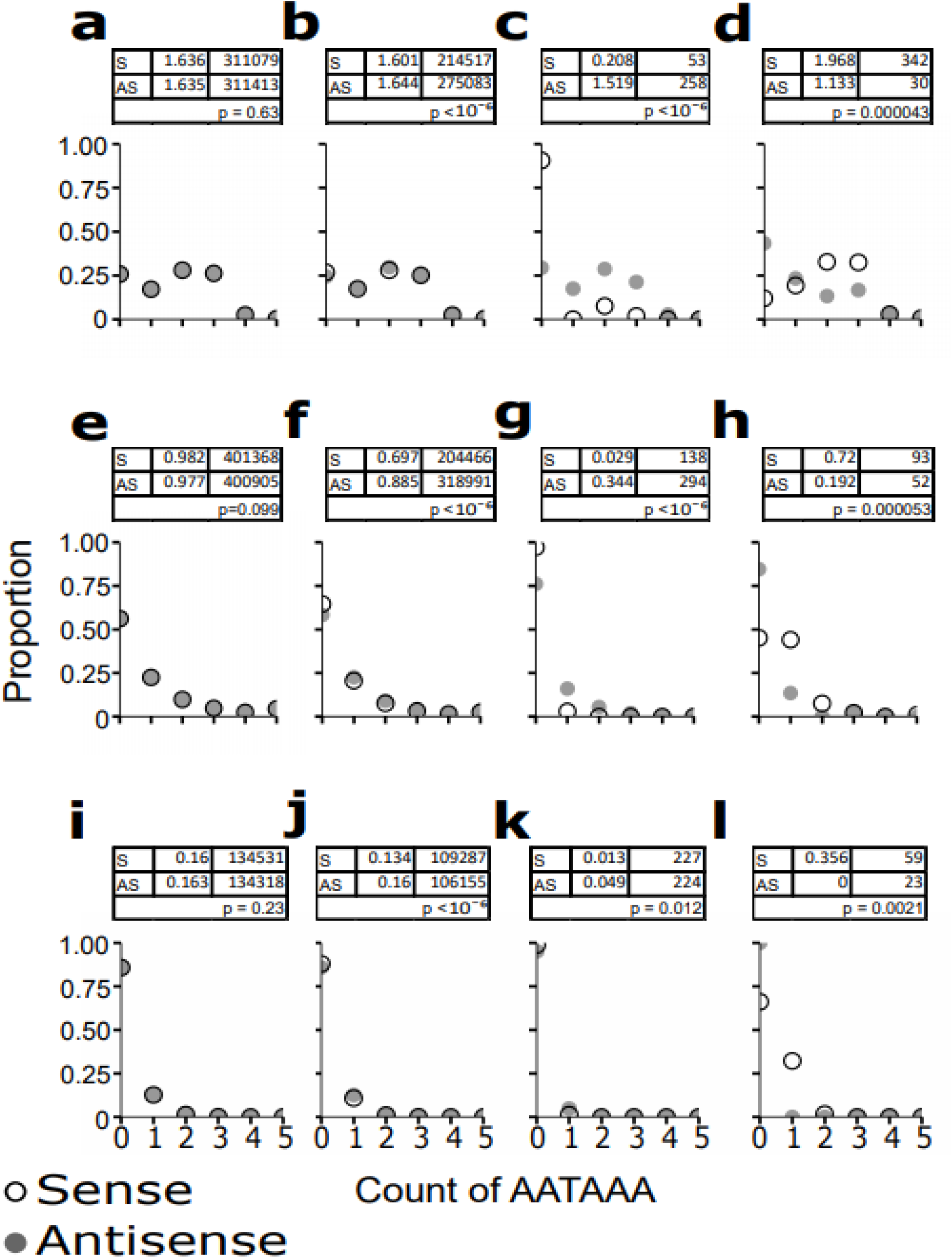
AATAAA PAS count significantly differs between sense and antisense-oriented retrotransposons in transcripts. In all cases the number of AATAAAs on the head-to-tail strand of the retrotransposon are counted. Each panel’s table summarizes sense (“S”; row 1) and antisense (“AS”; row 2)-oriented retrotransposons with the mean number of AATAAAs (column 2) and the total count of such elements in the dog reference genome CanFam3 (column 3). Row 3 reports the randomization test for difference in mean between S and AS (raw p value uncorrected for the 12 hypothesis tests). The rows of panels cover Can-SINEs (**a-d**), LINEs (**e-h**), and MIRs (**i-l**). The columns left to right are genomic locations: intergenic (**a, e, i**), intronic (**b, f, j**), within a 3’UTR but nearest edge >225 bp from the 3’ end (“far”; **c, g, k**), and within a 3’UTR but <225 bp from the 3’ end (“near”; **d, h, l**). For the intergenic location the “S” and “AS” denote top and bottom strand. In all histograms the count indicated at 5 is the number of copies with 5 or more AATAAAs.

Can-SINEs inserted within 3’UTRs far upstream from 3’ends are five times more likely to be in antisense than sense orientation. Furthermore, the antisense-oriented Can-SINEs have over a seven times higher mean count of AATAAA sequences (Fig. 5c). Most sense-oriented Can-SINEs in the “far” category have zero AATAAAs. The situation is just the reverse for “near” Can-SINEs, those wholly inserted within 225 bp of the 3’ end. In this location, 91% of Can-SINEs are inserted in sense orientation and they have significantly more AATAAAs than their antisense counterparts. A small number of sense-oriented Can-SINEs in the “near” location apparently have zero AATAAAs but after manually checking all such cases we found most are simply mis-annotated (see below).

LINEs on average have fewer AATAAAs in their sequence than do Can-SINEs but like Can-SINEs display a reversal in mean counts between the “near” vs. “far” categories (Fig. 5g, h). The ratios of numbers of LINE copies in sense vs. antisense are at most ~2-fold, much lower than the ~5-fold ratio for “far” Can-SINEs or the ~11-fold for “near” Can-SINEs. There are an equal number of MIRs inserted in sense and antisense orientation in the “far” region of 3’UTRs. The trend in the signal matches the Can-SINEs and LINEs in this location: sense-oriented MIRs have less than one-third the mean AATAAA count of antisense MIRs. However, the means are very low so the absolute number of MIRs with non-zero counts is also low and the difference is not significant after correction for multiple hypothesis testing (Fig. 5k). Only about 80 MIRs in the dog genome are in the “near” 3’UTR region and only a fraction of these (all in sense orientation) have an AATAAA. The pattern across all three types of retrotransposons is clear: the AATAAA count is elevated when a retrotransposon is inserted in sense orientation near the end of a 3’UTR. In contrast, AATAAA sequences occur at lower rates when the retrotransposon is farther upstream in a 3 ‘UTR or an intron.

We randomly sampled 20 3’UTRs in which a Can-SINE with one or more AATAAAs is “near” the 3’end in sense orientation. Twelve of these 3’UTRs do not occur in the orthologous gene in the human and eight do. Of the twelve, four occur in human ortholog introns, six occur downstream of the human ortholog and in two cases that are likely pseudogenes no human ortholog was identified. In the eight 3’UTRs with an orthologous human 3’UTR, four are longer in dog and four are shorter. For example, in the *DMAC2* gene, required for assembly of the mitochondrial complex I membrane arm [61], the dog 3’UTR of 592 bp terminates in a 13.8% sequence diverged sense-oriented SINEC_b1 with one AATAAA. This dog 3’UTR is supported by one unspliced expressed sequence tag but most of the 20 randomly sampled 3’UTRs are not.

Can-SINEs in sense orientation at the 3’ ends of 3’UTRs have two AATAAAs on average, nearly ten times the average for sense-oriented Can-SINEs farther upstream in 3’UTRs. We were therefore curious about the couple dozen Can-SINEs near the 3’end with a reported zero count of AATAAA. Perhaps these 3’UTRs have alternate PAS sequences in the SINE sequence. They may have a PAS in sequence unrelated to the SINE’s presence or do without a PAS. To distinguish among these possibilities, we individually checked each of these 38 SINE insertions (Table S3). All nine SINEC types of Can-SINE are represented in the set. In 32/39 (82%) of these 3’UTRs an AATAAA sequence is present within 40 bp of the 3’ end, so alternative PASs is not the primary explanation. In fact, in 26 of these 3’UTRs (strictly a subset of the 32 with near AATAAAs) the SINE insertion has been mis-annotated by being split into several separate repeat records. One of the records in each case is marked as simple sequence and spans an expansion of the TAAA(n) STR from the Can-SINE AT-rich tail. Therefore, in these 26/39 (66%) of cases the SINE insertion does include AATAAA, often in several copies.

The signal for counts of AATAAA imported by Can-SINEs to 3’ends is therefore even stronger than the distribution shown in Fig. 5d. Finally, there are seven 3’UTRs in the dog genome in which a sense-oriented Can-SINE is within our “near” category for 3’ends (meaning that all of the SINE insertion record is within 225 bp of the 3’ end) but the SINE end nearest the 3’UTR end is over 40 bp distant. Perhaps such locations are too far to productively provide an AATAAA in the SINE. Interestingly, in five of these 3’UTRs the SINE is missing most or all of its AT-rich tail. Possibly, deletions removing the PASs are favored when the SINE is this far upstream of the 3’ end.

One particularly interesting SINEC_Cf3 insertion occurred in dog *CTU1* (Fig. 6). *CTU1* along with *CTU2* kill melanoma cells [62]. The inactivation of the *CTU1-CTU2* complex causes loss of thiolation in tRNAs and is associated with ploidy abnormalities [63]. Essentially the entire 3’UTR modeled in the dog is the sense-oriented SINE. Although there are three separate repeat track records within this 3’UTR they belong to just a single Can-SINE insertion event, which is supported by the finding that 15 bp TSDs (with two mismatches) surround the three records. Therefore, the AATAAAs present in this 3’UTR were imported by the SINE. The small final coding exon and the SINE-become-3’UTR do not occur in the *CTU1* orthologs in two other carnivores: the ferret (*Mustela putorius furo*, reference musFur1) and cat (*Felis catus*, reference felCat9). In the reference genomes of both these species a trio of old repeat sequences, MIR, L2b and LTR106, are present in the middle of the intron, as in dogs, but the dog SINEC_Cf3 is not. Furthermore, the human (assembly hg38), ferret, and cat all show a completely different final coding exon and 3’UTR (Fig. 6a, c). Thus, a Can-SINE in one carnivore lineage, but not others, appears to have completely altered the downstream half of the *CTU1* gene.

**Fig. 6.**
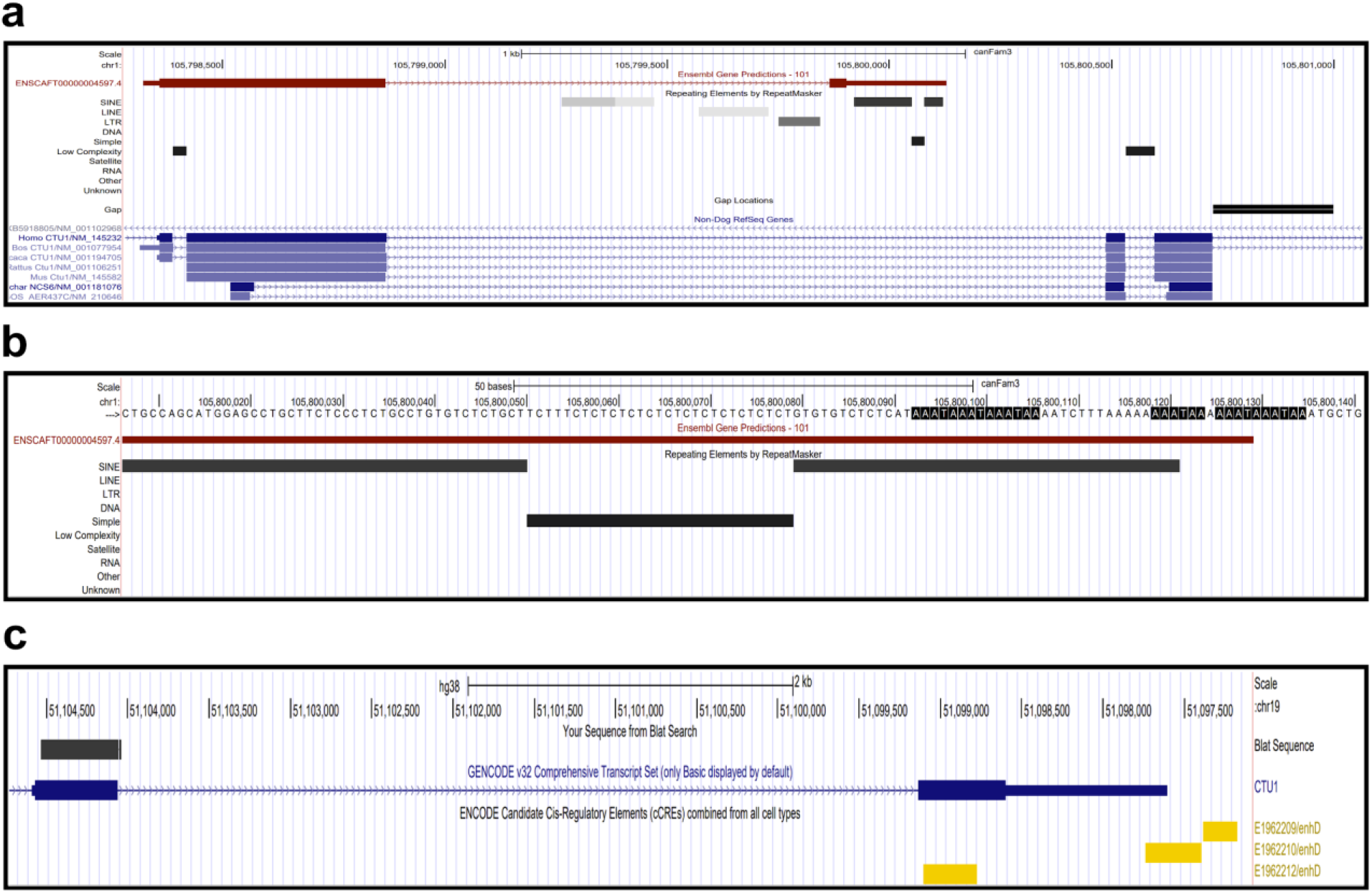
A sense-oriented SINEC_Cf3 is the dog *CTU1* 3’UTR. **a** The SINE is mis-annotated as three separate repeat records that are in fact one SINE insertion. **b** Six AATAAAs occur within the SINE’s AT-rich tail with a TCTTT RNA pol III terminator in the middle. A 15 bp target site duplication with two mismatches starts within the sixth AATAAA. **c** The human ortholog contains a different final coding exon and 3’UTR. The blatted sequence (rectangle on the left) indicates homology to the dog’s 2nd to last coding exon. The dog homolog of the human final coding exon can be seen in **a** in the non-dog track, blue.

A SINEC_Cf2 inserted into *SDHAF2* is another example of a mis-annotated sense-oriented Can-SINE near a 3’UTR 3’ end (Fig. S3). In mice with *SDH5* knocked out, epithelial cells in the lungs have elevated mesenchymal markers, causing EMT [65]. Again, the entire 3’UTR in the dog gene model is the SINE, although mis-annotation has broken it into two separate records. But the 3’UTR’s AATAAA is followed by the TCTTT RNA pol III terminator motif and then a polyA run are the AT-rich tail of the SINE (Fig. S3b). This is supported by a 15 bp TSD with one mismatch that starts in the last four As of the homopolymer run. The matching TSD is the end of the protein-coding exon. This SINEC_Cf2 element is relatively young at just 4.3% sequence divergence from the consensus sequence and is not present in either the ferret or cat orthologs of *SDHAF2*. There are several variant 3’UTRs for the human ortholog, some which are much longer than what is present in the dog (Fig. S3c). Possibly the dog gene is missing some 3’UTR regulation as a result of this SINE insertion.

The human *IL17RA* gene’s 3’UTR is over 5800 bp long but in the dog ortholog a Can-SINE insertion has apparently truncated nearly all of the 3’UTR (Fig. S4). *IL17RA* signaling has a role in the pathogenesis of psoriatic arthritis in both the human and mouse [66]. This SINEC_Cf2, at 12.8% sequence divergence from consensus, is likely older than many other insertions but it nevertheless is not present in either the ferret or cat *IL17RA* orthologs. It would be interesting to check for this SINE insertion in other Canidae family members.

The young SINEC_Cf copy in sense orientation at the 3’end of the dog *TTLL12* 3’UTR (Fig. S5) is just 4.4% diverged from the SINEC_Cf consensus sequence and is not present in either the ferret or cat orthologs. As in the examples above, the single SINE insertion has been annotated as multiple records. The two records should be annotated as a single SINE insertion, however, as shown by the perfect 16 bp TSDs that surround the two segments. Strikingly, the AT-rich tail of this SINE has mutated into a pure TAAA(17) simple tandem repeat, providing many AATAAA sequences in the process. TTLL12 acts as an inhibitor of the activation of cellular antiviral signaling [67].

The high copy number of Can-SINEs in the dog genome implies that short TAAA(n) simple tandem repeats are delivered to many thousands of loci by retrotransposon activity. STR mutation rates are often orders of magnitude higher than single base substitutions and in some cases the TAAA(n) experiences a repeat expansion, as seen in the dog *TTLL12* locus (Fig. S5). Perhaps in part due to Can-SINE activity, the dog has a high genome count of TAAA(n) STRs compared to other vertebrates (Table S4), with roughly twice as many of these STRs as are found in the human, chimpanzee, or mouse genomes.

Our distribution of AATAAA sequences in 3’UTRs is a bit surprising (Fig. 1). The rounded peak with mode ~22-24 bp was expected but the single-base spikes at 28, 32, and 36 bp were not. In order to confirm that our Python code was accurately enumerating the AATAAAs at the same distances as are found in the reference genome, we randomly selected 50 3’UTR records from each of the top five peak bp positions for AATAAA frequency: 22, 24, 28, 32, and 36 bp from the 3’UTR 3’ end. We manually checked each of these sampled loci and verified that an AATAAA does occur at that bp position (Table 1; Table S5). We count the distance to the 3’UTR 3’ end from the promoter-proximal 5’ end of all motifs. For each of these 250 3’UTRs we also recorded if there was an AATAAA found at any of the other four positions (Table S5). We found that Can-SINEs contribute to the 4 bp interval peaks: 24, 28, 32, and 36. In fact, Can-SINEs contributed the AATAAA at 60% of our 32 bp sample and in every case the 3’UTR also has an AATAAA 4 bp upstream. Thus, part of the explanation for the 36 bp spike may be this correlation with AATAAA presence at the 32 bp position. However, that alone can’t explain all of the 36 bp spike since it is nearly five times greater magnitude than the 32 bp spike. Given the TAAA(n) repeat in the Can-SINE AT-rich tail, the 4 bp interval is understandable.

**Table 1.** A random sample of 250 3’UTRs having AATAAA at each of the five peak frequencies (50 UTRs sampled per distance). In parenthesis is the number that also have AATAAA at 36 bp.

|  | AATAAA Identified at (bp) |  |  |  |  |
| --- | --- | --- | --- | --- | --- |
|  | 22 | 24 | 28 | 32 | 36 |
| Can-SINE | 0 (0) | 7 (4) | 4 (3) | 31 (31) | 6 (6) |
| LINE | 3 (0) | 4 (0) | 2 (2) | 0 (0) | 5 (5) |
| MIR | 0 (0) | 0 (0) | 0 (0) | 0 (0) | 1 (1) |
| Other | 47 (0) | 39 (1) | 44 (7) | 19 (3) | 38 (38) |

We next tabulated the absolute counts of SINEs in sense and antisense orientation that contribute AATAAA to the 3’ end of 3’UTRs (Table S6). Many times more AATAAAs are imported to 3’UTR ends by sense-oriented rather than antisense-oriented Can-SINEs. This pattern does not hold for MIRs where the numbers are more equal. But very few AATAAAs in 3’ ends are found within MIRs at all. The 4 bp periodicity in the Can-SINE AATAAAs is evident for the 28, 32, and 36 bp positions for each of the Can-SINE sub-types except SINEC_c1 and perhaps SINEC_c2.

## Discussion

Can-SINEs have been active throughout the evolution of the dog species, including in recent time. The youngest Can-SINE, SINEC_Cf, appears to still be active today as evidenced by the tens of thousands of polymorphic insertions in purebred dogs [25]. In contrast, other types like SINEC_c1 and c2, are much older. We have annotated individual insertions of Can-SINE elements to better understand and predict the ages of various copies. Length of polyA tail has been used as a predictor for age of SINE insertions [59]. Insertions of non-A may “stabilize” the polyA sequence. We find a correlation between the length distribution of perfect TSDs and SINEC type (Fig. 4), where the younger SINEC_Cf, Cf2 and Cf3 types exist in many more copies with perfect 15 bp TSDs. SINEC_Cf copies with perfect 15 bp TSDs also have on average longer AT-rich tails. We found a number of specific SINE insertions in which the AT-rich tail underwent an expansion in the TAAA(n) STR, including in cases where the Can-SINE is providing the PAS AATAAA (or set of AATAAAs) for a 3’UTR. It would be interesting to know the timing of this process. Are these STRs more mutable when the SINE is freshly inserted and less likely to have inclusions of sequence within the tandem repeat [68]? Perhaps an optimum time window exists after a SINE inserts but before it accumulates many mutations in which it is more likely to provide a PAS to a gene.

Why do we find Can-SINEs in 3’UTR ends that have an expanded TAAA(n) STR? Is this a chance occurrence resulting simply from the high mutability of the STR? Or are these cases of an adaptive strengthening of the PAS (if indeed additional AATAAA sequences do strengthen the termination signal)? Possibly, added AATAAAs serve as a “work-around” for sub-optimal sequences surrounding the PAS? We have identified several hundred sense-oriented Can-SINEs at the 3’ends of 3’UTRs and the dataset includes all nine of the sub-types from old SINEC_C1s to young SINEC_Cfs. Nearly all of them contain AATAAAs. This suggests a long-lasting and ongoing evolutionary process of SINE-PAS importation to gene ends.

With hundreds of domestic dogs now being whole genome sequenced, and more and more Canidae family members getting a first individual sequenced, the data exist to compare orthologous Can-SINEs across canids and analyze the extent to which gene usage of Can-SINE PASs may be conserved. To what extent have Can-SINE insertions driven switching to new polyadenylation sites in canid evolution? Perhaps through comparative genomics, too, signatures of selection may reveal adaptive Can-SINE-altered gene ends.

We speculate that Can-SINE insertion-caused shortening of 3’UTRs (Fig. S4) could expose or remove miRNA binding sites and lead to changes in gene expression, as has been demonstrated in humans and other mammals [69]. Changes in 3’UTR length can bring miRNA binding sites closer to the ends of the 3’UTR where they are more effective [70]. In human *CDC6*, for example, 3’UTR shortening removes negative regulation of gene expression [71].

The high rate of recent SINEC_Cf insertions in the domestic dog genome provides a natural experiment for assessing the strength of transcription termination signals in Can-SINEs. We previously identified tens of thousands of putative polymorphic SINEC_Cf elements in a set of purebred dogs, including insertions within 3’UTRs. If these data were paired with gene expression datasets it would be possible to experimentally check use or non-use of Can-SINE-PASs in 3’UTRs and elsewhere in gene transcripts. Presumably, switching to a newly inserted Can-SINE PAS is a disruptive event for a gene and likely to be under strong negative selection. We expect such SINE insertions to be rare alleles and our previous evidence supports that [25].

Our analysis here is limited in several ways. First, we analyze only the SINE insertions present in the Boxer reference genome. There are many more insertion loci in the dog population generally. Second, the gene models are likely to have limitations too. Coding exons tend to get good annotation coverage earlier than untranslated regions [72]. More complete annotation of 3’UTRs as additional gene expression datasets are reported will certainly help. One of our findings in particular is a surprise that may be related to gene modeling. In analyses of 3’UTR 3’ ends in, for example, Beaudoing et al [18], a single mode peak of high frequency AATAAA occurrence is evident in the dataset. We find exactly the same peak in the same place. Our numbering system is from 5’ ends of the hexamers but when translated 6 bp matches Beaudoing et al [18]. However, overlaid on our peak are spikes of single base peaks at four bp intervals. The largest of these spikes by far is at precisely 36 bp from 3’UTR 3’ ends. Fearing bugs in our bioinformatics, we manually checked 250 randomly picked 3’UTRs and found AATAAAs exactly matching our Python script distance calls in every case. The four bp interval of the spikes is at least partly explained by the TAAA(n) periodicity in Can-SINE AT-rich tails (Fig. S6), which causes AATAAA to recur every four bp. Less clear, however, is why the 36 bp distance in particular should be so favored. Genome-scale experiments focused on mapping 3’ends of dog genes will be needed to resolve this question.

Can-SINEs are a T+ category SINE with putative transcript termination potential. Previously, we found that in dog, as in other mammals, SINEs in introns strongly favor the antisense orientation [25], consistent with the idea that sense-orientated Can-SINEs are selected against, presumably due to a potential to disrupt gene expression. Here we’ve provided additional support for the idea that Can-SINEs have RNA pol II termination potential because Can-SINEs in transcripts have significant differences in AATAAA counts when in sense vs. antisense orientation (Fig. 5a-d). The magnitude of this difference is large in upstream portions of 3’UTRs but small (though significant) in introns. We find the same signal for insertions of LINEs and MIRs in introns too. Within 3’UTR upstream regions the mean AATAAA count in Can-SINEs drops to 0.2, far lower than the 1.5 average AATAAAs present on antisense Can-SINEs in the same gene region. Thus, AATAAA removal via mutation appears to be favored in this region. Many Can-SINEs in sense orientation in 3’UTR upstream regions have zero AATAAAs. It would be worthwhile to investigate with expression datasets whether the Can-SINEs there that do have AATAAAs can in fact serve as polyadenylation signals in dog cells and if so, how frequently this occurs. Orthologs for at least some of these Can-SINEs likely exist in closely related canid species and a check could then be made for correlation between the presence or absence of AATAAAs in those species vs. polyadenylation signal usage in the Can-SINE.

As more reference genomes are collected and annotated with gene models, it will be interesting to compare our results in canids with other T+ SINEs in other mammal clades. A dog Can-SINE newly inserting into an intron or 3’UTR very likely has more intrinsic transcription termination potential than human *Alu*, but perhaps not more than other T+ SINEs like rabbit CSINEs or mouse B2s [38]. We predict that other T+ SINE classes, when analyzed, will show a similar signal of AATAAA suppression for sense-oriented SINEs in transcripts.

The genomic era has seen an explosive increase in our knowledge of SINE and LINE retrotransposons and their roles in the evolution of genes and genomes. One theme that emerges is the specificity and particularity with which each lineage has been impacted by these “genome pests”. Primate *Alu* has been shown to serve as a PAS [57] but first requires activating mutations that convert the sequence into a PAS. Sauria SINEs found in reptiles from geckoes to snakes to *Anolis carolinensis* similarly appear to lack potential for transcription termination [73]. In contrast, the T+ SINEs in diverse mammal groups may be playing significant roles over evolutionary time in shaping the ends of gene transcripts. As we collect reference genomes from ever greater numbers of species in diverse lineages, we’re getting to read these particular stories of genomes being shaped by many thousands of SINEs and other repeated sequences.

## Conclusions

Here we find that dog Can-SINEs, like other T+ SINEs, carry multiple copies of the polyadenylation signal motif AATAAA in their AT-rich 3’ end tails. Can-SINEs inserted within introns and 3’UTRs have significantly different mean AATAAA counts depending on their strand and we conclude that AATAAA is not a neutral sequence in either context. We find the same pattern for both LINEs and MIRs in introns and also for LINEs in 3’UTRs. Furthermore, we find that many more sense-oriented Can-SINEs are inserted in 3’UTR 3’ ends than into positions farther upstream in 3’UTRs. We conclude that Can-SINE retrotransposition has imported AATAAA PASs into genes and in many loci these have then been used to signal transcription termination. For several loci we find evidence that a Can-SINE insertion has truncated a 3’UTR or created a new one. We hypothesize that other carnivores, also carrying Can-SINEs at high copy numbers, have experienced the same kinds of evolutionary changes.

## Authors’ contributions

J.D.C. analyzed most of the datasets and produced most figures. N.B.S. led the study, conducted the randomization tests, and wrote the Python programs for DNA segment intersection analysis, motif finding, and SINE annotation. L.A.D. analyzed all SINE annotations, produced Table S4, and helped produce most figures. J.D.C. and N.B.S. planned all analyses and wrote the paper with input from L.A.D.

## Acknowledgements

We thank Jonathan Specht for help with functions within our SINE annotation script.

