## Supplement for "SINE Retrotransposons Import Polyadenylation Signals to 3’UTRs in Dog (*Canis familiaris*)"

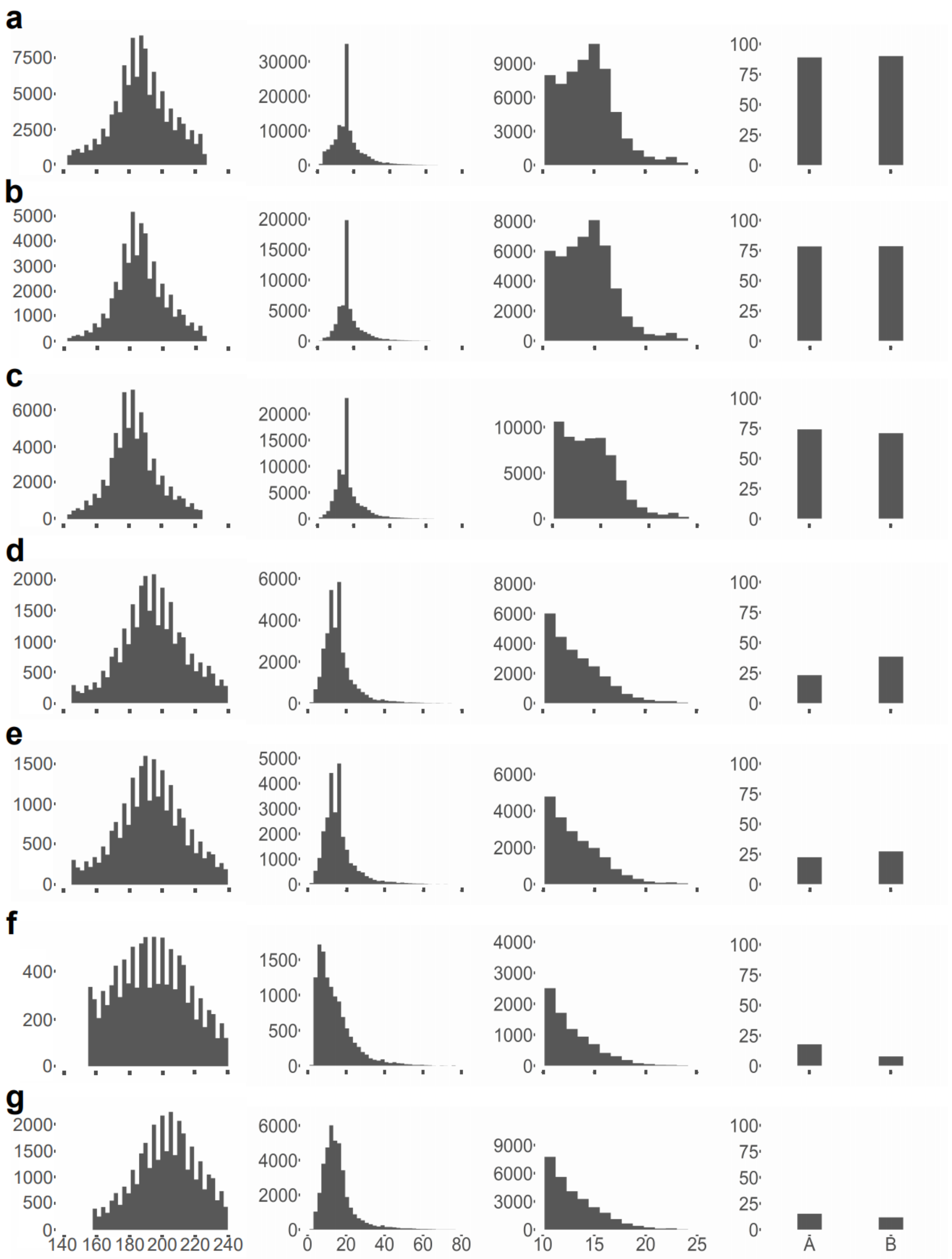


**Fig. S1** Annotation of Can-SINE elements from the dog reference genome. Column 1 is length histograms of the sequence identified between TSDs, column 2 is a length histogram for the AT-rich tail sequence, and column 3 is histograms of perfect TSD lengths. All histograms have horizontal axis bp and vertical axis count. Column 4 shows the percentage of the SINEs in which A box and B box sequences were identified. **a** SINEC_Cf2, **b** SINEC_Cf3, **c** SINEC_a2, **d** SINEC_b1, **e** SINEC_b2, **f** SINEC_c1, **g** SINEC_c2.

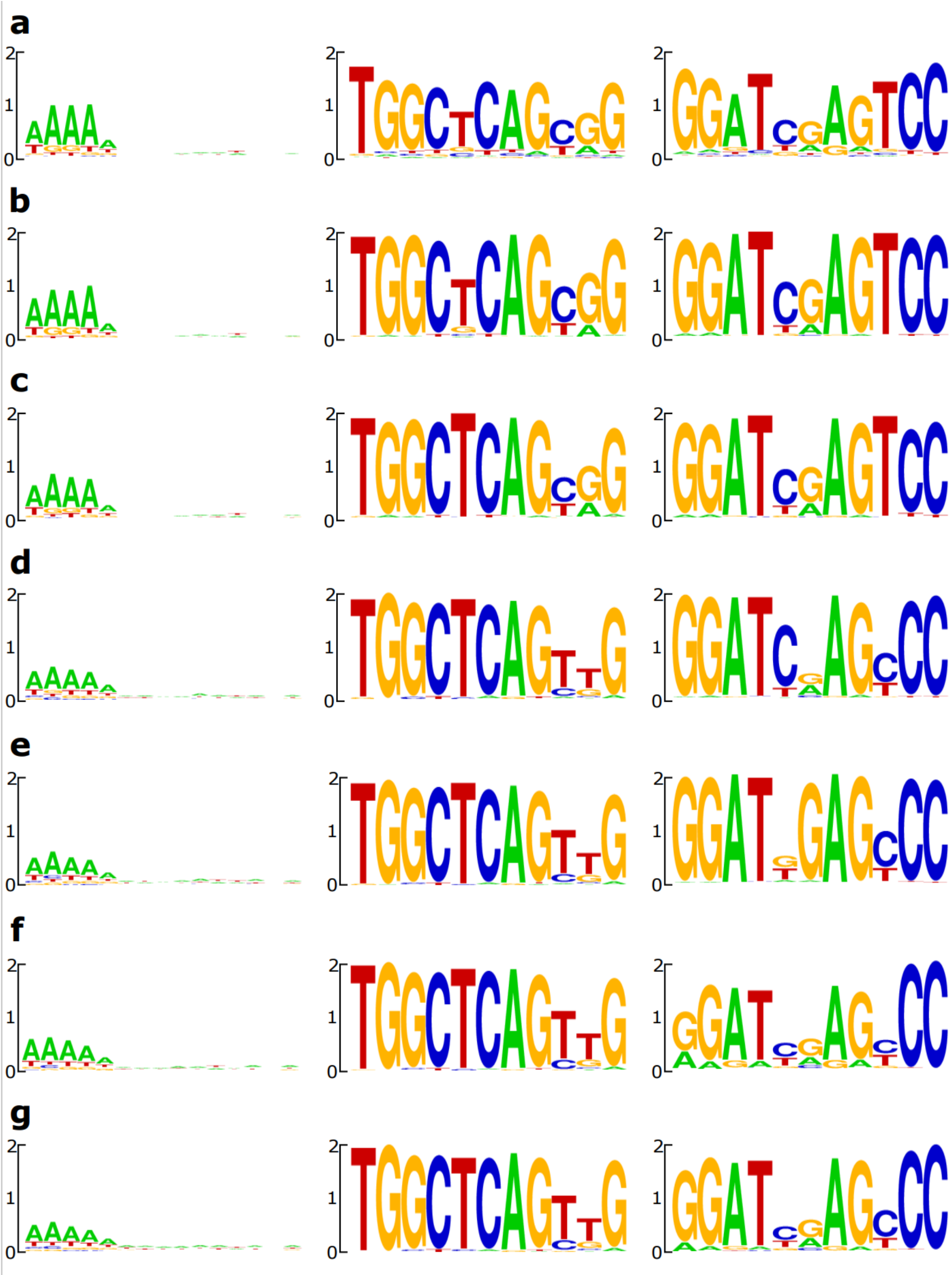

**Fig. S2** Sequence logos of Can-SINE types from the dog reference genome. Columns left to right are of sequence logos of the TSD, A box, and B box. **a** SINEC_Cf2, **b** SINEC_Cf3, **c** SINEC_a2, **d** SINEC_b1, **e** SINEC_b2, **f** SINEC_c1, **g** SINEC_c2.

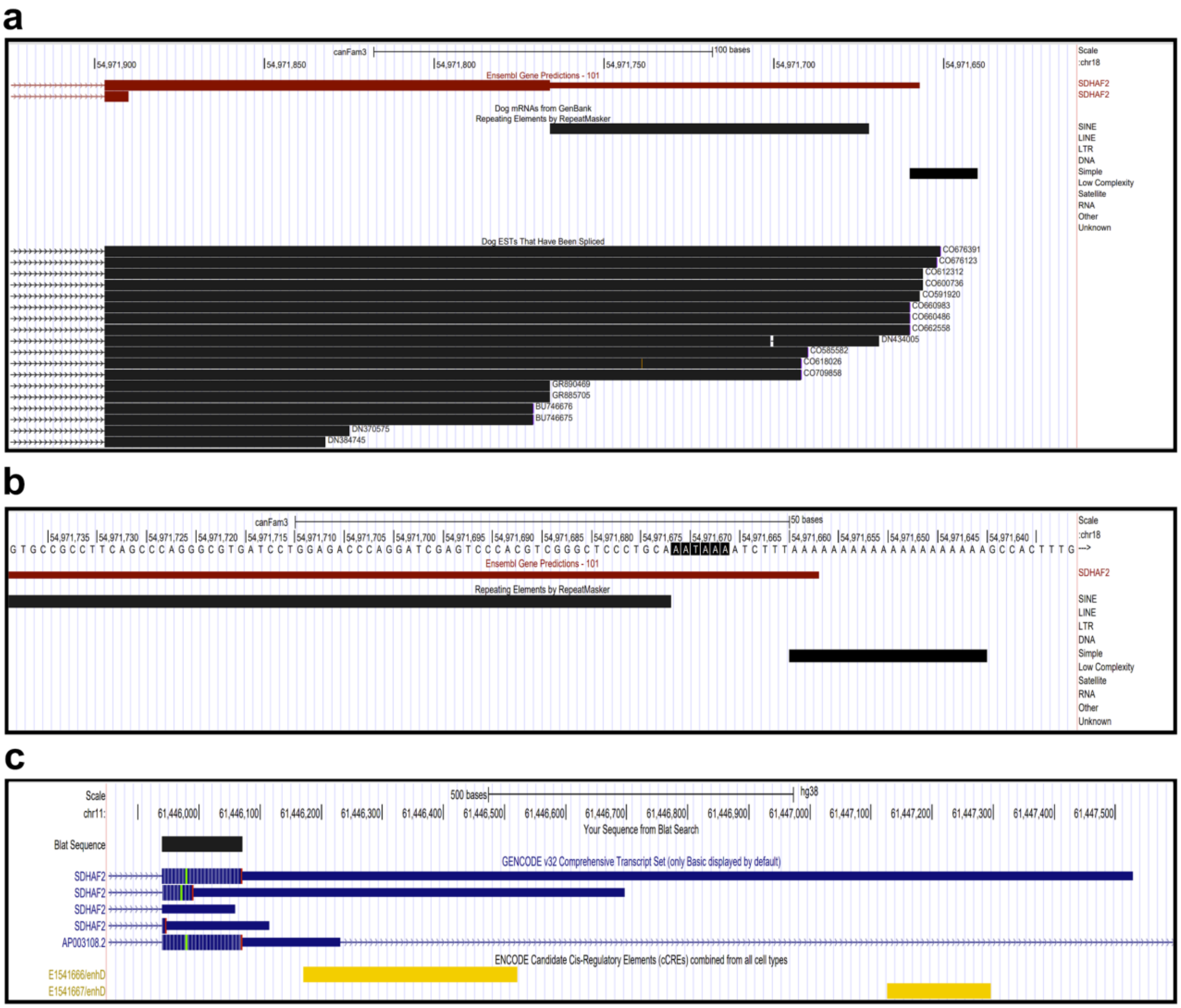


**Fig. S3** A sense-oriented SINEC_Cf2 is the entire 3'UTR for the *SDHAF2* gene. **a** dog spliced expressed sequence tags terminate in the modeled 3'UTR. **b** The SINE is annotated as separate SINE and simple sequence records but the SINE record should continue through the AATAAA, the TCTTT RNA pol III terminator motif, and the polyA. Starting with the last four As of the homopolymer run a 15 bp TSD with one mismatching base matches the end of the coding exon immediately upstream of the SINE's head end. **c** The human ortholog's 3'UTRs are variable and include cis-regulatory elements.

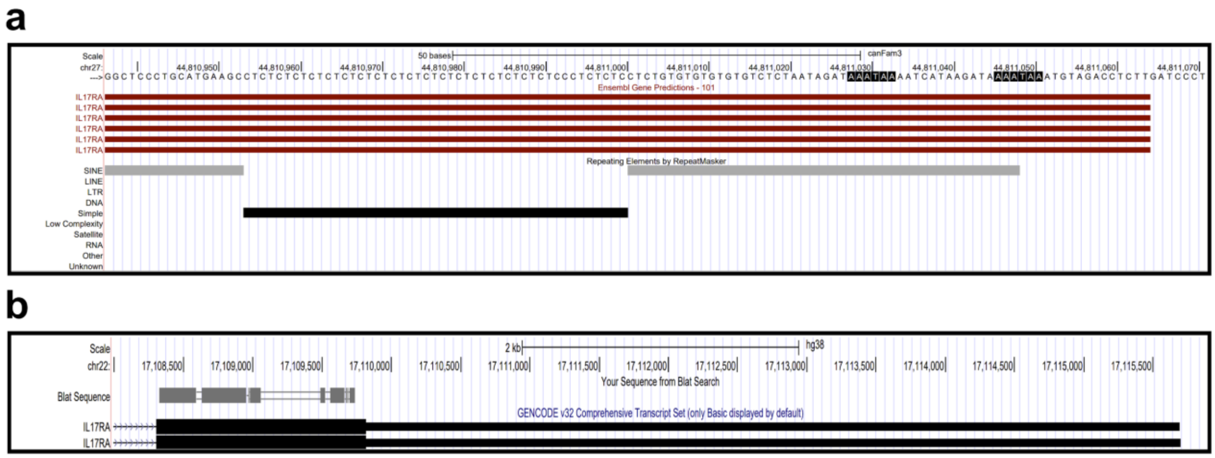


**Fig. S4** A sense-oriented SINEC_Cf2 is over half of the ~320 bp dog 3’UTR of *IL17RA*. **a** The single SINE insertion is annotated as three separate records: a head portion, a CT(n) simple repeat, and the AT-rich tail in which AATAAA motifs are found at 19 and 36 bp from the 3’ end of the 3’UTR. A 13 bp TSD starts at the TAAA of the 3’-most AATAAA. **b** In the human ortholog the 3’UTR is >5800 bp long. The Blat Sequence track shows homology between the last coding exon in the dog and the human.

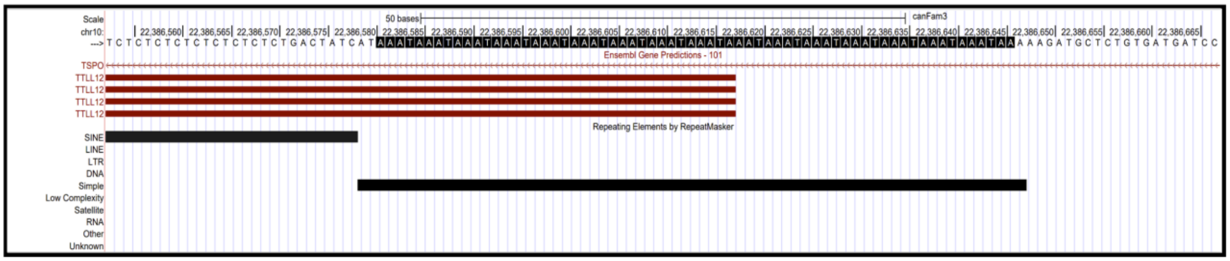


**Fig. S5** A sense-oriented young SINEC_Cf is the source of an uninterrupted TAAA(17) STR expansion providing many AATAAA motifs at the 3’ end of the dog *TTLL12* 3’UTR. A 16 bp perfect target site duplication starts immediately after the STR.

### SUPPLEMENTARY TABLES

**Table S1** Genome-wide counts of 11 PAS motifs in the 3’ ends of 3’UTRs.

**Table S2** Counts of 11 PAS motifs in Can-SINE and MIR consensus sequences.

**Table S3** Can-SINEs in sense orientation within 225 bp of 3’UTR 3’ ends that have apparently zero AATAAA motifs are mostly mis-annotated.

**Table S4** Genome-wide counts in several mammals and other vertebrates of simple tandem repeats of the form {AAAT, AATA, ATAA, TAAA, ATTT, TATT, TTAT, TTTA}.

**Table S5** Confirmation that our intersections python script correctly identifies AATAAA sequences within 3’UTRs. We manually checked 50 random 3’UTR for each of five AATAAA distances.

**Table S6** Genome-wide counts of AATAAAs within retrotransposons near the 3’ end of 3’UTRs. They mostly occur in sense-oriented elements at 4-bp intervals 28, 32, and 36 bp from the 3’ end.
